# CLIC4 is regulated by RhoA-mDia2 signaling through Profilin-1 binding to modulate filopodium length

**DOI:** 10.1101/259259

**Authors:** Elisabetta Argenzio, Katarzyna M. Kedziora, Leila Nahidiazar, Tadamoto Isogai, Anastassis Perrakis, Kees Jalink, Wouter H. Moolenaar, Metello Innocenti

## Abstract

CLIC4 is a cytosolic protein implicated in diverse actin-based processes, including integrin trafficking, cell adhesion and tubulogenesis. CLIC4 is rapidly recruited to the plasma membrane by G_12/13_-coupled receptor agonists and then partly co-localizes with β1 integrins. Receptor-mediated CLIC4 translocation depends on actin polymerization, but the mechanism and functional significance of CLIC4 trafficking are unknown. Here we show that RhoA activation by either LPA or EGF is necessary and sufficient for CLIC4 translocation, with a regulatory role for the RhoA effector mDia2, an inducer of actin polymerization. We find that CLIC4 directly interacts with the G-actin-binding protein Profilin-1 via conserved residues that are required for CLIC4 trafficking and lie in a concave surface. Consistently, silencing of Profilin-1 impaired CLIC4 trafficking induced by either LPA or EGF. CLIC4 knockdown promoted the formation of long integrin-dependent filopodia, a phenotype rescued by wild-type CLIC4 but not by trafficking-incompetent CLIC4(C35A). Our results establish CLIC4 as a Profilin-1-binding protein and suggest that CLIC4 translocation provides a feedback mechanism to modulate mDia2/Profilin-1-driven cortical actin assembly and membrane protrusion.

## Introduction

Chloride intracellular channel (CLIC) proteins (CLIC1–6) are small globular proteins (~28 kDa) that are structurally related to the omega-class glutathione S-transferases (GSTOs), showing an N-terminal thioredoxin-like domain followed by an all α-helical C-terminal domain [1–4]. However, CLICs have distinct but poorly understood cellular functions from the GSTs. Contrary to their original name, CLIC proteins do not function as conventional chloride channels but instead have been implicated in diverse actin-dependent processes, such as tubulogenesis, membrane remodeling, endosomal trafficking, vacuole formation and cell adhesion, among others [3–5]. It has recently been shown that CLICs have intrinsic glutaredoxin-like activity, at least under *in vitro* conditions with a conserved reactive cysteine serving as key catalytic residue [6,7], but whether CLIC glutaredoxin-like activity is maintained in the reducing cytosol is unknown. Moreover, there is no convincing evidence that CLICs may act as intracellular redox sensors.

CLIC4 is arguably one of the best studied CLIC family members. Despite decades of research, progress in CLIC function has been frustratingly slow, partly because direct binding partners have been elusive. CLICs are often found associated with the cortical actin cytoskeleton and are detected on intracellular membranes, where they may participate in the formation and maintenance of vesicular compartments [5,8–11]. Growing evidence indicates that CLIC proteins play roles in actin-mediated trafficking events. CLIC4 knockout mice are viable but are smaller and show defects in actin-dependent processes, including delayed wound healing and impaired endothelial and epithelial tubulogenesis [12–14]. Strikingly, CLIC4 undergoes rapid redistribution from the cytosol to the plasma membrane in response to G_12/13_-coupled receptor agonists, notably lysophosphatidic acid (LPA, a major serum constituent) and other GPCR agonists [15,16]. CLIC4 translocation was strictly dependent on RhoA-mediated actin polymerization and, interestingly, on the reactive but enigmatic Cys35 residue as well as on other conserved residues that in GSTs are critical for substrate binding [15]. This strongly suggests that the substrate-binding features of the omega GSTs have been conserved in the CLICs, along with the fold itself, and that binding of an as yet unknown partner (or substrate) is essential for CLIC4 function. Yet the functional relevance of agonist-induced CLIC4 trafficking remains obscure to date.

In epithelial cells, CLIC4 is homogeneously distributed and can colocalize with a subset of early and recycling endosomes [10]. In response to serum or LPA stimulation, CLIC4 rapidly colocalizes with β1 integrins, consistent with CLIC4 functioning in actin-dependent exocytic-endocytic trafficking under the control of receptor agonists [15]. A recent study on renal tubulogenesis confirmed that CLIC4 regulates intracellular trafficking, showing that CLIC4 colocalizes with the retromer complex and recycling endosomes, while CLIC4 depletion resulted in the enrichment of branched actin at early endosomes [13]. Collectively, these findings establish CLIC4 as a trafficking regulator that acts in concert with the actin cytoskeleton.

A major challenge towards a better understanding of the CLICs is the identification of specific binding partners; this should help to clarify how CLICs traffic to or associate with membrane compartments. In the present study, we characterize CLIC4 trafficking and function in further mechanistic detail and establish the G-actin-binding protein Profilin-1 as a direct binding partner of CLIC4. Our results indicate that, through Profilin-1 binding, CLIC4 functions in a RhoA-mDia2 signaling network to integrate actin dynamics, membrane protrusion and cell adhesion.

## Results and Discussion

### Rapid but transient translocation of CLIC4 to the plasma membrane induced by LPA and EGF

In serum-deprived neuronal and epithelial cells, CLIC4 resides mainly in the cytosol, where it is highly mobile [15], and to a low extent in distinct patches at the plasma membrane. Using HeLa cells, we found that CLIC4 is rapidly recruited to the plasma membrane not only by G_12/13_-RhoA-coupled receptor agonists such as LPA, but somewhat unexpectedly also by a prototypic receptor tyrosine kinase ligand, notably epidermal growth factor (EGF) (**Fig. 1A, and Suppl. Movies 1 and 2**). Receptor-mediated CLIC4 accumulation at the plasma membrane coincided with CLIC4 depletion from the cytosol (**Fig. 1B** and **1D**), occurring within seconds of receptor stimulation. Maximum membrane recruitment was reached within ~1 min of LPA or EGF addition. Subsequently, CLIC4 gradually disappeared from the plasma membrane as it translocated back to the cytosol over a time course of about 10 min. in the continuous presence of agonist (**Fig. 1B-E**). Similar to LPA, EGF induced CLIC4 to translocate to β1 integrins at the plasma membrane ([10] and EA, unpulbished results). CLIC4 translocation induced by either LPA or EGF was blocked by the G-actin-binding toxin Latrunculin A, indicating that CLIC4 trafficking depends on F-actin polymerization ([15]. Furthermore, using an EGFR inhibitor, we ruled out that LPA acts through EGFR transactivation. It thus appears that EGF and G_12/13_-coupled receptors share a common signaling mechanism to evoke CLIC4 trafficking.

**Figure 1.**
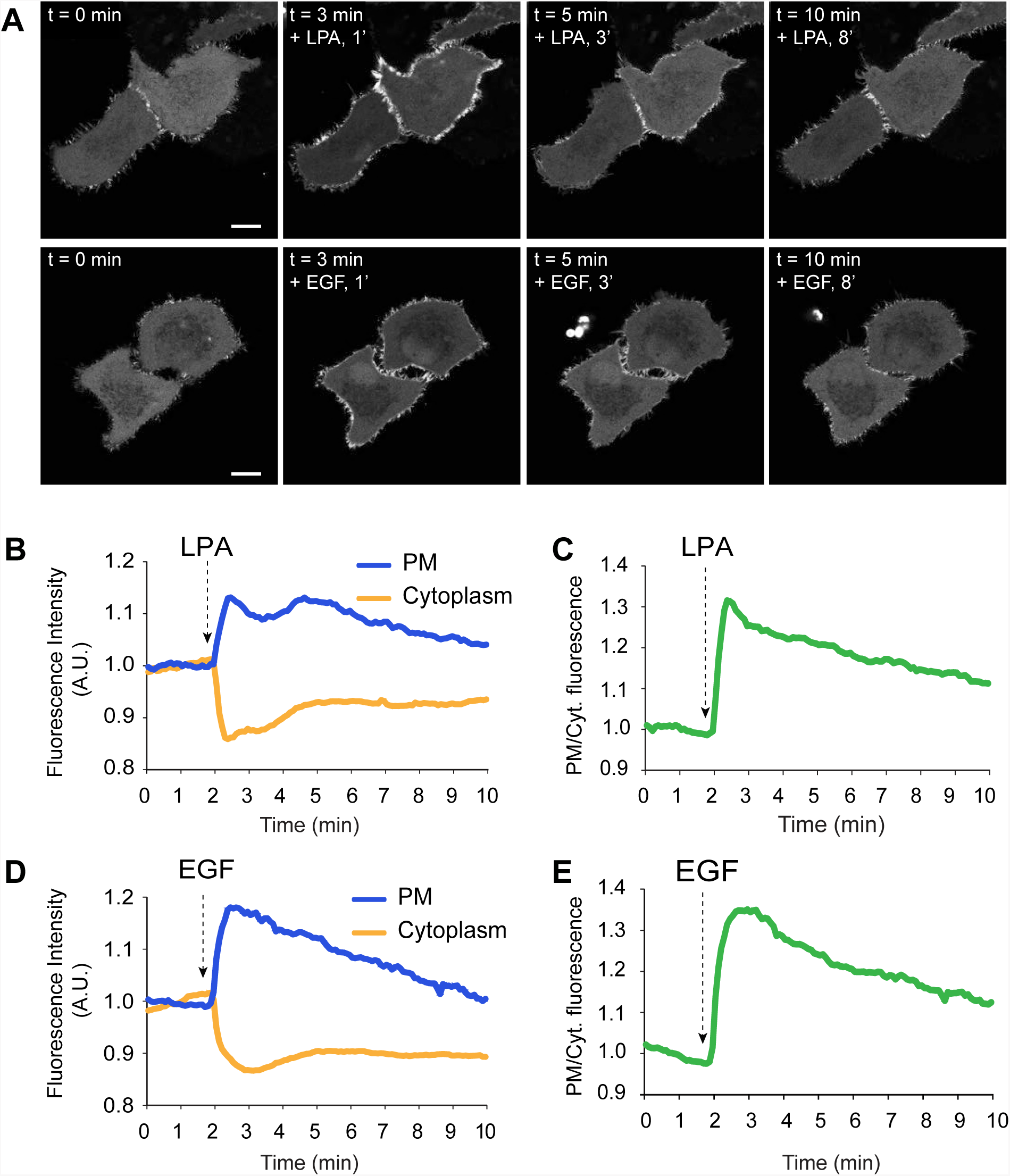
LPA- and EGF-induced translocation of CLIC4 to the plasma membrane. (**A**) Live-cell imaging of CLIC4 translocation to the plasma membrane. Cells were seeded on glass coverslips and transfected with YFP-CLIC4. LPA (2 µM, top) and EGF (100 ng/ml) were added 2 minutes after starting imaging. Frames from time-lapse movies at the indicated time points are shown. Scale bar: 10 µm. (**B-E**) Quantification of CLIC4 translocation. LPA- (B) and EGF-induced (D) translocation was quantified by measuring YFP fluorescence at the plasma membrane (PM, blue line) and in cytoplasm (Cyt, yellow line). (**C,E**) Net translocation is expressed as PM/Cyt fluorescence ratio (LPA, n= 18 cells; EGF n= 14 cells from two independent experiments).

CLIC4 shows oxireductase activity towards artificial substrates *in vitro,* with Cys35 serving as active-site residue, which is inhibited by ethacrynic acid and IAA-94, compounds once used as chloride channel blockers [6]. Since CLIC4 trafficking depends on residue Cys35 [15], we pretreated the cells with these membrane-permeable inhibitors but found no effect on agonist-induced CLIC4 trafficking (EA, unpublished results). These results argue against intrinsic oxireductase activity playing a role in CLIC4 trafficking.

### RhoA activation is necessary and sufficient to drive CLIC4 translocation

Given the apparent involvement of RhoA in CLIC4 translocation [15], we set out to monitor RhoA activation in real-time in LPA- and EGF-stimulated cells, as well as in CLIC4-depleted cells, using a RhoA-specific FRET-based biosensor [17]. As shown in **Fig. 2A**, both LPA and EGF rapidly activated RhoA in HeLa cells. Somewhat unexpectedly, EGF was a stronger RhoA activator than LPA in these cells. Peak activation of RhoA by either LPA or EGF was reached within one minute; thereafter, Rho-GTP levels gradually decreased over a 10 min. time period, stabilizing at a level above pre-stimulation values. CLIC4 knockdown (using two independent shRNAs) consistently led to a small increase in basal RhoA-GTP levels as revealed in pull-down assays (**Fig. 2B-C**), possibly secondary to altered integrin function. However, CLIC4 depletion did not affect agonist-induced RhoA activation, neither in amplitude nor in kinetics (**Fig. 2A-B**). It is of note that the RhoA activation kinetics closely parallels the time course of CLIC4 translocation (**Fig. 2A**), consistent with RhoA activity providing the driving force for CLIC4 recruitment to the plasma membrane.

**Figure 2.**
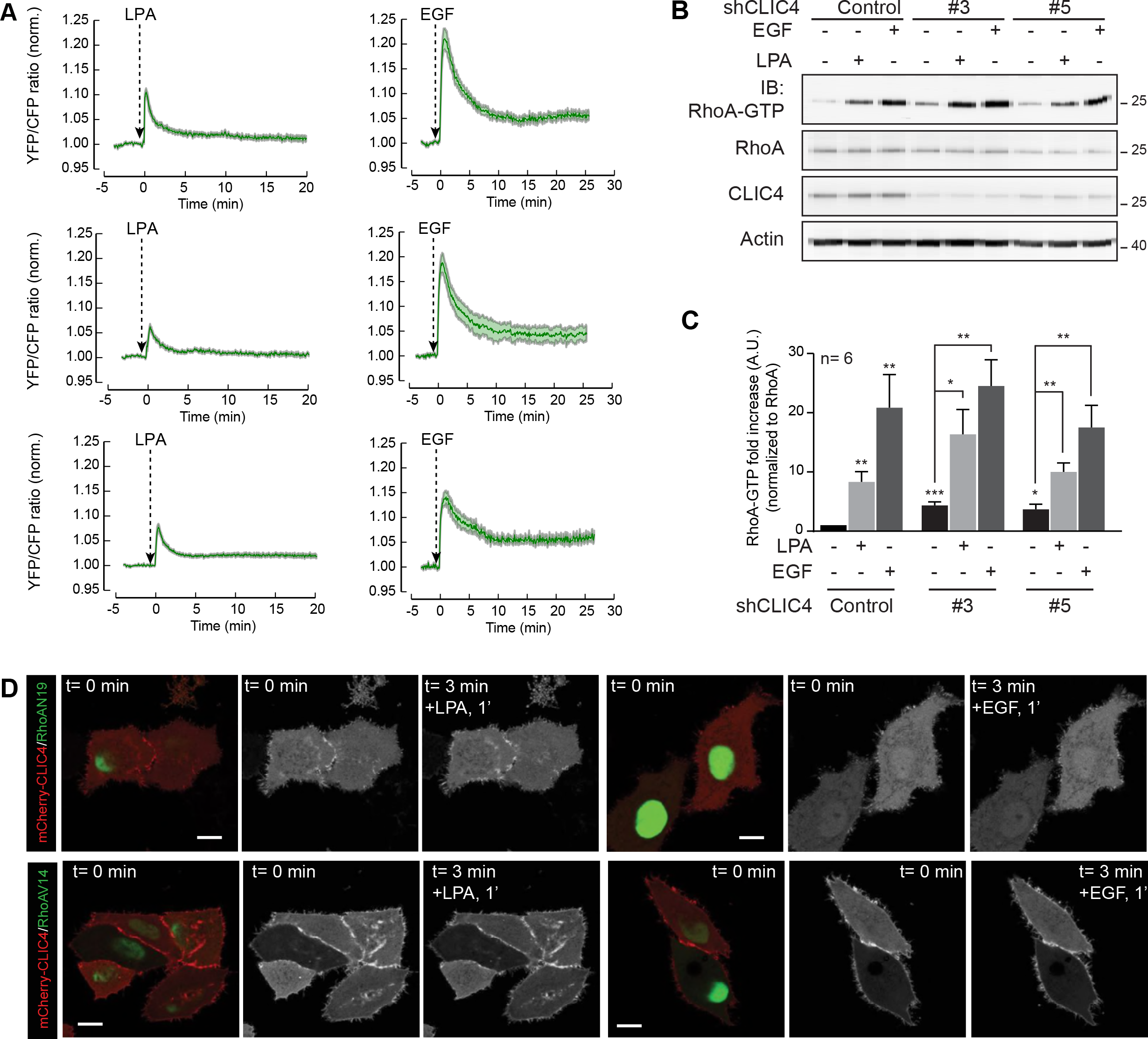
LPA- and EGF-induced translocation of CLIC4 depends on RhoA activation. (**A**) Kinetics of RhoA activation by LPA and EGF and dependency on CLIC4. shControl and shCLIC4 knockdown cells were transfected with a RhoA biosensor [17]. RhoA activity is plotted as normalized YFP/CFP ratio over time. (**B,C**) RhoA pull-down assays. shControl and shCLIC4 knockdown HeLa cells were serum starved overnight and either stimulated with LPA (5 µM) or EGF (100 ng/ml) for 3 minutes or left untreated. GTP-bound RhoA was pulled down as described in Materials and Methods. GTP-bound and total RhoA were detected by immunoblot analysis using anti-RhoA antibodies. CLIC4 knockdown was monitored by immunoblot analysis of total cell lysates anti-CLIC4 antibody. Actin was used as loading control. Representative blots of one out of six independent experiments are shown. Densitometric analysis (mean + s.e.m.) of six experiments is shown in (**C**). (**D**)HeLa cells were plated on glass coverslips and co-transfected with mCherry-CLIC4 and either dominant negative (RhoAN19, top) or constitutively active RhoA (RhoAV14, bottom) in a bicistronic IRES vector expressing GFP. Cells were serum starved and either stimulated with LPA (2 µM) or EGF (100 ng/ml). RhoAN19 and RhoAV14 transfected cells express monomeric GFP. Frames from time-lapse movies at the indicated time points are shown. Scale bar: 10 µm.

Indeed, expression of dominant-negative RhoA(N19) blocked agonist-induced CLIC4 translocation (**Fig. 2D**), while constitutively active RhoA(V14A) forced CLIC4 to accumulate permanently at the plasma membrane (**Fig. 2D**). Under the latter conditions, translocation of the remaining cytosolic CLIC4 could not be further stimulated by LPA (**Fig. 2D**). Thus, RhoA activation is a necessary and sufficient signal for CLIC4 to translocate. Moreover, these results imply that agonist-induced CLIC4 translocation to the plasma membrane may serve as a convenient readout of RhoA activation.

### Involvement of Rho effector mDia2

Major downstream effectors of RhoA are Rho-kinase (ROCK) and the mDia formins. ROCK induces actomyosin-mediated cell contraction, while the formins mDia1 and mDia2 promote Rho GTPase-regulated nucleation and elongation of linear actin filaments to drive cell protrusion. ROCK inhibition did not affect CLIC4 translocation (**Suppl. Fig. 1** and [15]). We therefore focused our attention on mDia1 and mDia2, which are the two major formins in HeLa cells [18].

Stable knockdown of either mDia1 or mDia2 (using lentiviral vectors shmDia1 and shmDia2, respectively) did not alter the subcellular localization of CLIC4 (**Fig. 3A,C**) nor its total expression level (**Fig. 3E**). While CLIC4 translocation was not affected by mDia1 knockdown, it was prominently reduced upon mDia2 depletion (**Fig. 3A-D**; second independent hairpin shown **in Suppl. Fig. 2**). Moreover, the pan-Formin inhibitor SMIFH2 [19,20] caused a similar decrease in CLIC4 translocation (**Fig. 3A-D**). Thus, translocation of CLIC4 is in large part regulated by the RhoA-mDia2 signaling axis through cortical actin polymerization. However, we found no interaction between recombinant CLIC4 and purified F-actin in co-sedimentation experiments. Since CLIC4 recruitment was not fully inhibited upon mDia2 knockdown, it is noteworthy that both LPA and EGF also induced activation of Rho-GTPase family member Cdc42 (**Fig. 3F**). However, LPA activates Cdc42 through G_i_-signaling [17], whereas CLIC4 trafficking is independent of G_i_ [15]. We therefore conclude that agonist-induced CLIC4 translocation is regulated, at least in large part, by RhoA-mDia2-driven actin polymerization at the cell periphery.

**Figure 3.**
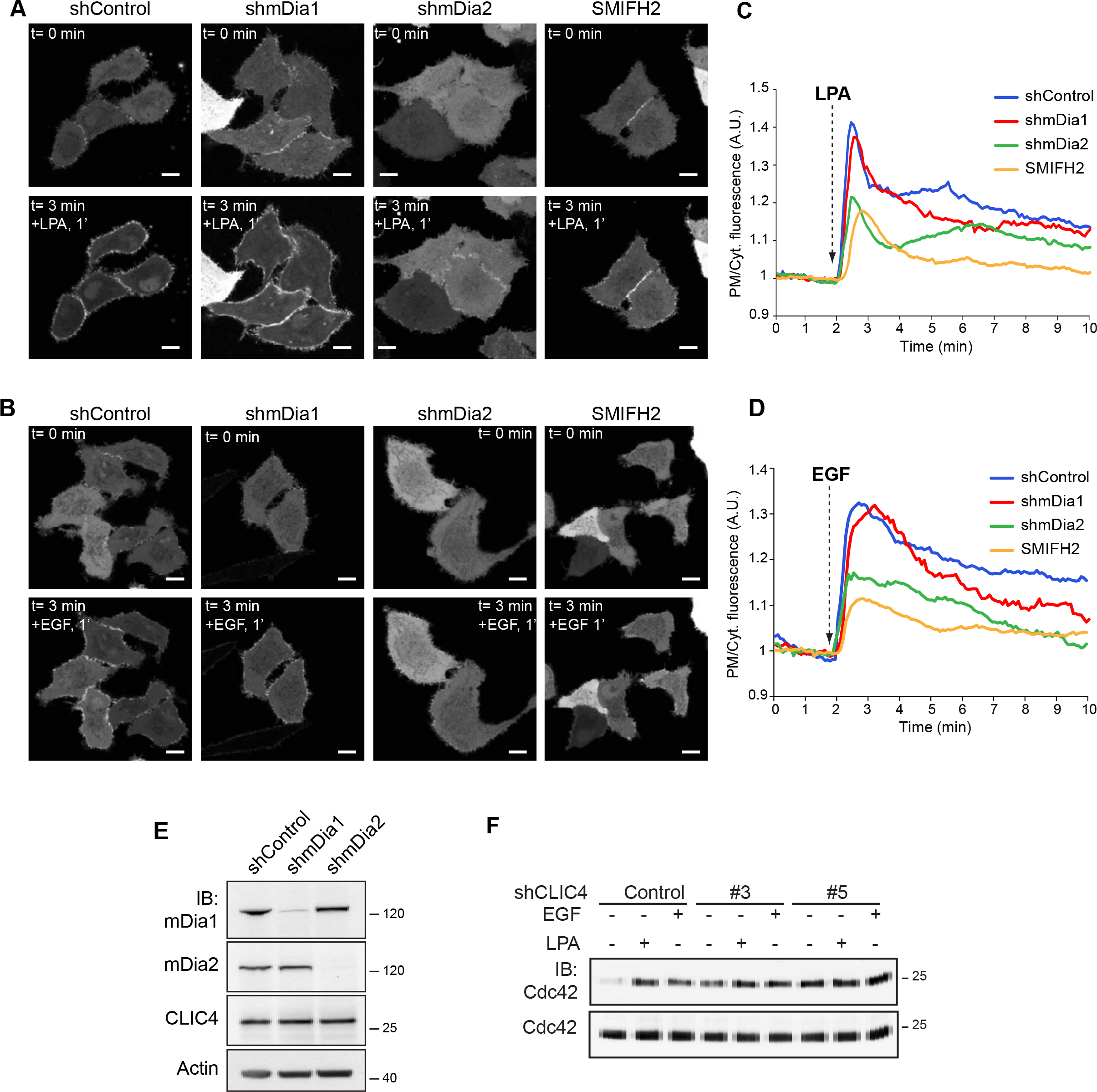
LPA-induced translocation of CLIC4 relies on mDia2 and its activity. (**A,B**) Live-cell imaging of CLIC4 translocation in mDia knockdown HeLa cells. Stable mDia1 and mDia2 knockdown cells were obtained as described in the Materials and Methods. shControl, shmDia1 and shmDia2 cells were seeded on glass coverslips and transfected with YFP-CLIC4. shControl cells were pre-treated with SMIFH2 (50 µM, 20 min) or left untreated. LPA (**A**) (2 µM) or EGF (**B**) (100 ng/ml) were added 2 minutes after starting imaging. SMIFH2 was maintained during stimulation. Frames from time-lapse movies at the indicated time points are shown. Scale bar: 10 µm. (**C**) Quantification of LPA-induced translocation. shControl (blue trace, n= 9), shmDia1 (red trace, n= 9), shmDia2 (green trace, n= 13) and SMIFH2-treated cells (yellow trace, n= 14), from two independent experiments. (**D**) Quantification of EGF-induced translocation. shControl (blue trace, n= 10), shmDia1 (red trace, n= 9), shmDia2 (green trace, n= 9) and SMIFH2-treated cells (yellow trace, n= 9), from two independent experiments. Net translocation is (C-D) is expressed as PM/Cytoplasm ratio. (**E**) mDia knockdown efficiency was determined by immunoblotting (IB) using anti-mDia1 and anti-mDia2 antibodies. β-actin was used as loading control. (**F**) Cdc42 pull down. shControl and shCLIC4 knockdown HeLa cells were serum starved overnight and either stimulated with LPA (5 µM) or EGF (100 ng/ml) for 3 minutes or left untreated. GTP-bound Cdc42 was pulled down as described in Material and Methods. GTP-bound and total Cdc42 were detected by immunoblot analysis using anti Cdc42 antibody. Representative blots of one out of three independent experiments are shown.

### CLIC4 binds Profilin-1 via conserved residues, including Cys35

Since CLIC4 translocation depends on mDia2 activity, we focused our attention on the mDia2-interacting protein Profilin-1 (encoded by *PFN1),* which modulates the activity of formins [18,21,22]. Profilin-1 is a ubiquitous G-actin-binding protein with separate binding sites for actin and poly-Pro stretches, such as those found in the FH1 domains of formins [22,23]. We examined a possible interaction, both biochemical and functional, between CLIC4 and Profilin-1. As shown in **Fig. 4A-B**, Profilin-1 co-immunoprecipitated with CLIC4 in transfected HEK293 cells. However, the translocation-incompetent CLIC(C35A) mutant failed to interact with Profilin-1 (**Fig. 4A-B**). We established that the interaction is direct by using recombinant purified CLIC4 and Profilin-1 in pull-down assays (**Fig. 4C-E**). Remarkably, direct binding of Profilin-1 was impaired not only in CLIC(C35A) but also in other CLIC4 mutants that fail to translocate [15], notably the F37D, P76A, D97A and Y244A mutants (**Fig. 4E**). These respective residues are highly conserved and lie in a concave surface (or “cleft”) of CLIC4, similar to that observed in the omega GSTs where it mediates glutathione binding [1,15]. However, CLIC proteins exhibit only very low affinity for glutathione. Yet it is conceivable that CLICs might use this cleft as a binding site for an extended macromolecular chain, notably a polypeptide or a post-translationally modified protein, to be targeted to a particular subcellular location [1,3,15,24,25].

**Figure 4.**
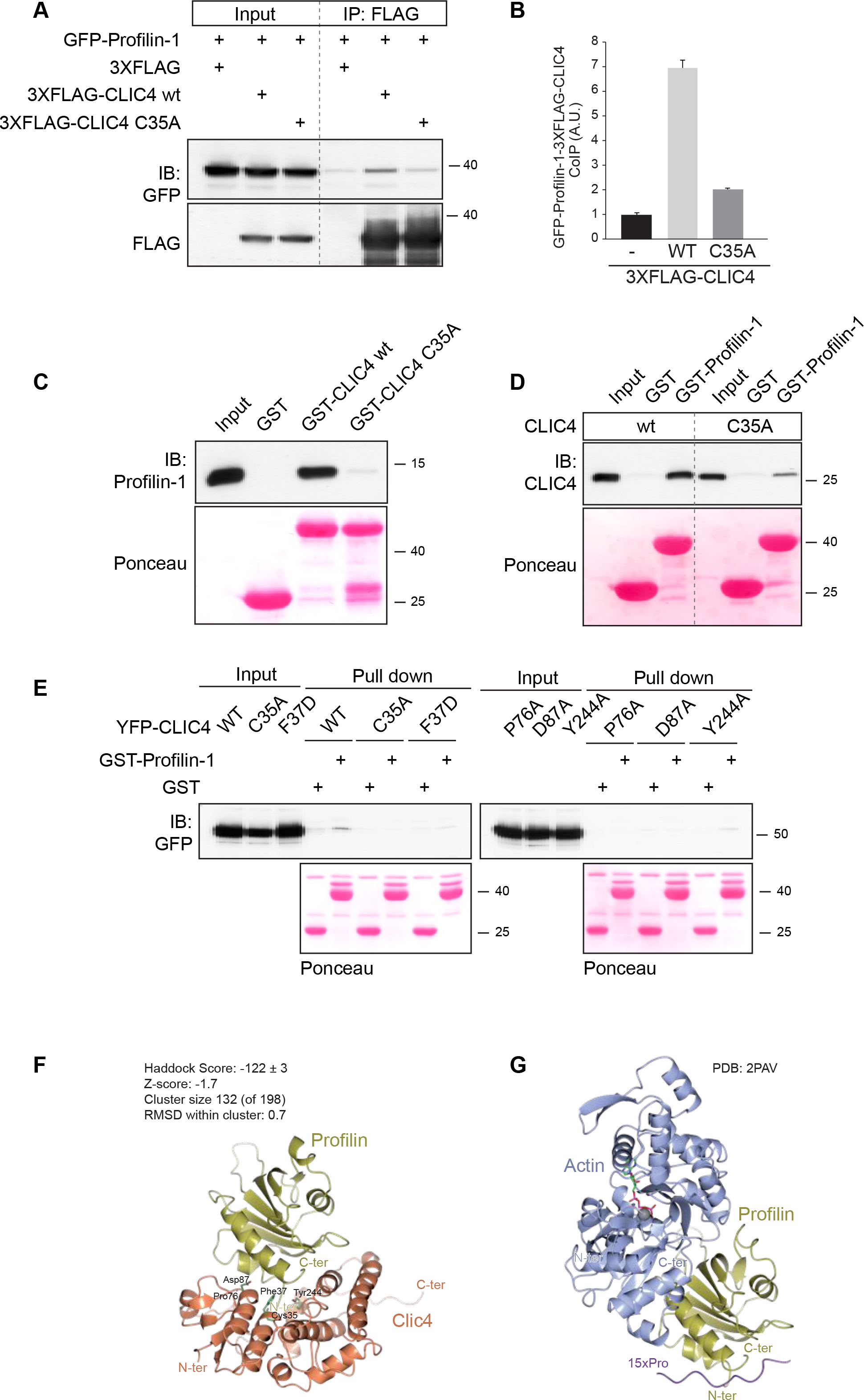
CLIC4 binds Profilin1 via distinct residues that make up an open cleft. (**A**) Profilin-1 coimmunoprecitates with CLIC4 in cells. 3XFLAG-CLIC4 wild type (wt) and its mutant in Cys35 (C35A) were cotransfected with GFP-PFN1 in HEK cells. CLIC4 was immunoprecipitated from cell lysate (1 mg) using anti FLAG antibody. Coimmunoprecipitated Profilin-1 and CLIC4 were detected by immunoblotting using anti-GFP and anti-FLAG antibodies, respectively. Representative blots of one out of three independent experiments are shown. (**B**) Densitometric analysis (mean + SD) of three experiments is shown. (C-D) CLIC4 directly binds Profilin-1 *in vitro.* (**C**) Purified immobilized GST-CLIC4 wt and GST-CLIC4 C35A mutant were incubated with Profilin-1 for 2 hrs on ice and pulled down with GST-Agarose beads. The amount of Profilin-1 pulled down by GST-CLIC4 was detected by immunoblotting using anti-Profilin-1 antibodies. The reciprocal experiment was performed using GST-Profilin1 and CLIC4 wt and the C35A mutant (**D**). GST alone was used as a control. Ponceau staining showed equal loading of the GST-fusion proteins. (**E**) YFP-CLIC4 (wt) and the indicated mutants were transfected into HEK293 cells. Total cell lysate (1 mg) was incubated with GST or GST-Profilin-1. The amount of YFP-CLIC4 wt and mutants pulled down by Profilin-1 was detected by immunoblot analysis using anti-GFP antibody. Ponceau staining showed equal loading of the GST-fusion proteins. Representative blots of one out of two independent experiments are shown. (**F,G**) Molecular modeling of Profilin-1 interactions. (**F**) HADDOCK computational model showing CLIC4 (orange) and Profilin-1 (gold). The CLIC4 residues discussed in the Results, and the N- and C- termini of both proteins are indicated. (**G**) The crystal structure of Profilin-1 (gold) binding to G-actin (blue) (PDB:2PAV). See text for further details.

To gain insight into the CLIC4-binding site of Profilin-1, we examined whether CLIC4 may regulate the actin-binding properties of Profilin-1. To this end, we exploited that Profilin-1 inhibits actin self-assembly by forming a 1:1 complex with G-actin. We found no effect of recombinant CLIC4 on the kinetics of Profilin-1-actin polymerization measured *in vitro* (**Suppl. Fig. 3A bottom**), showing that CLIC4 and G-actin do not compete for Profilin-1. Moreover, CLIC4 did not affect spontaneous actin polymerization (**Suppl. Fig. 3A top**) or bind to F-actin (results not shown). Together, these results show that CLIC4 in itself has no direct effects on actin dynamics.

### Molecular modeling of the CLIC4-Profilin-1 complex

We set out to create a model of the CLIC4-Profilin-1 complex using the High Ambiguity Driven protein-protein DOCKing (HADDOCK) modeling suite [26,27]. HADDOCK modelling resulted in a large cluster (132 models, 67% of total) of similar models (0.7Å RMSD from the overall lowest-energy structure) with excellent scores (**Fig. 4F**). The CLIC4 interaction surface is located on the other end of the Profilin-1 binding site to G-actin, and is similar to the binding region of poly-Proline (poly-Pro) peptides (**Fig. 4G**) [28,29]. Thus, the modelling results corroborate our biochemical observations that CLIC4 and G-actin do not compete for Profilin-1 binding with a plausible structural explanation.

In an attempt to determine the equilibrium dissociation constant of the CLIC4-Profilin-1 complex, we used surface plasmon resonance (SPR) but could only detect weak direct binding that did not reach saturation (estimated K_D_>200 µM; A. Fish and T. Heidebrecht, unpublished results). This may not come as a surprise, since physiologically relevant interactions mediated by the poly-Pro binding site of Profilin-1 are in a similar range [28,30]. Moreover, as we previously pointed out, weak and/or dynamic interactions may allow CLICs to transiently interact with distinct binding partners along a given trafficking route [3]. In any case, these results establish Profilin-1 as a direct binding partner of CLIC4, and they link Profilin-1 binding to the capability of CLIC4 to respond to RhoA activation.

### Involvement of Profilin-1 in LPA- and EGF-induced CLIC4 translocation

To examine the importance of Profilin-1 in CLIC4 trafficking, we generated Profilin-1-depleted HeLa cells using two distinct shRNAs; these cells showed unaltered CLIC4 expression and localization (**Fig. 5A-B**). LPA- and EGF-induced CLIC4 translocation to the plasma membrane was strongly reduced upon Profilin-1 depletion (**Fig. 5B-C**). Furthermore, the formation of filopodia induced by mDia2 was virtually abolished in these cells (**Suppl. Fig. 3B**). These results support the view that Profilin-1 binding connects CLIC4 to RhoA-mDia2 regulated actin dynamics.

**Figure 5.**
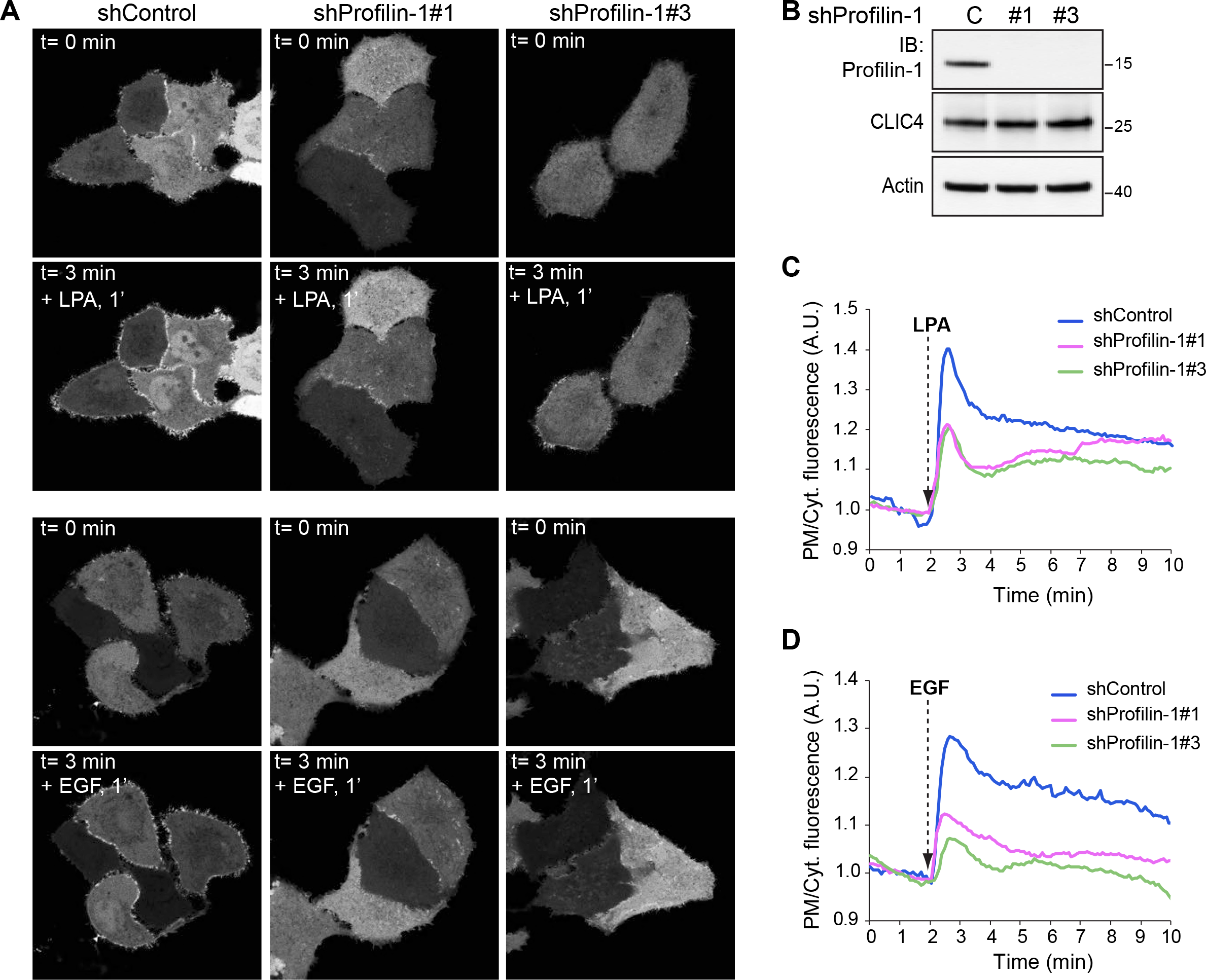
Role of Profilin-1 in LPA-induced CLIC4 translocation. (**A**) Live-cell imaging of CLIC4 translocation in Profilin-1 knockdown HeLa cells. Stable knockdown cells were obtained as described in the Materials and Methods. Two distinct shRNAs targeting Profilin-1 are shown. (**B**) Knockdown efficiency as determined by immunoblotting (IB) using anti-Profilin-1 antibody; β-actin was used as loading control. Cells were seeded on glass coverslips and transfected with YFP-CLIC4. LPA (2 µM) and EGF (100 ng/ml) were added 2 minutes after starting imaging. Frames from time-lapse movies at the indicated time points are shown. Scale bar: 10 µm. (**C**) Quantification of LPA-induced CLIC4 translocation (n_shControl_= 13, n_shPFN1#1_= 13, n _shPFN1#3_= 19 cells, from two independent experiments). (**D**) Quantification of EGF-induced CLIC4 translocation (n_shcontrol_= 12, n_shPFN1#1_= 13, n_shPFN1#3_= 20 cells, from two indepedent experiments). Net translocation in (**C, D**) is expressed as the PM/Cyt ratio.

### CLIC4 does not affect mDia2-driven actin polymerization in vitro

The similarities between the poly-Pro and CLIC4 binding sites of Profilin-1 suggests that CLIC4 may regulate mDia2-driven actin dynamics. Since Profilin-1 accelerates formin-mediated actin assembly [31,32], we measured actin polymerization induced by the FH1-FH2 domain of mDia2 in the presence of Profilin-1 with or without recombinant CLIC4. mDia2 increased the rate of actin polymerization, as expected, but this was not affected by CLIC4 (**Suppl. Fig 3A bottom**). We ruled out that CLIC4 interferes directly with mDia2 by repeating these assays without Profilin-1 (**Suppl. Fig. 3A, top**). Thus, CLIC4 does not affect mDia2 activity *in vitro* when actin polymerization occurs in solution. Since the binding affinities of CLIC4 and mDia2 for Profilin-1 are both the high μM range, it remains possible that CLIC4 and mDia2 may compete for Profilin-1 at membranes where bi-dimensionality increases their local concentration.

### CLIC4 depletion promotes integrin-dependent filopodium protrusion

In addition to altering integrin trafficking [10], we found that CLIC4 depletion triggered the formation of long dynamic filopodium-like protrusions (**Fig. 6A-B**). These protrusions were positive for the F-actin-bundling protein Fascin (**Fig. 6C**) and showed mDia2 expression at the tips (**Fig. 6D**). Filopodium formation is regulated by formins and can be induced by Cdc42 GTPase activity [33–35]. Consistent with this, Cdc42 activity was upregulated upon CLIC4 knockdown (**see Fig. 3F**) while the formin inhibitor SMIFH2 caused these protrusions to disappear (**Fig. 6E**). The observed finger-like protrusions therefore qualify as genuine filopodia. Given the effects of CLIC4 knockdown on integrin signaling and cell adhesion [10], these filopodia may reflect an integrin-dependent phenotype. Indeed, filopodia were similar when control and CLIC4 knockdown cells were grown on poly-lysine, which implies dependency on integrins.

**Figure 6.**
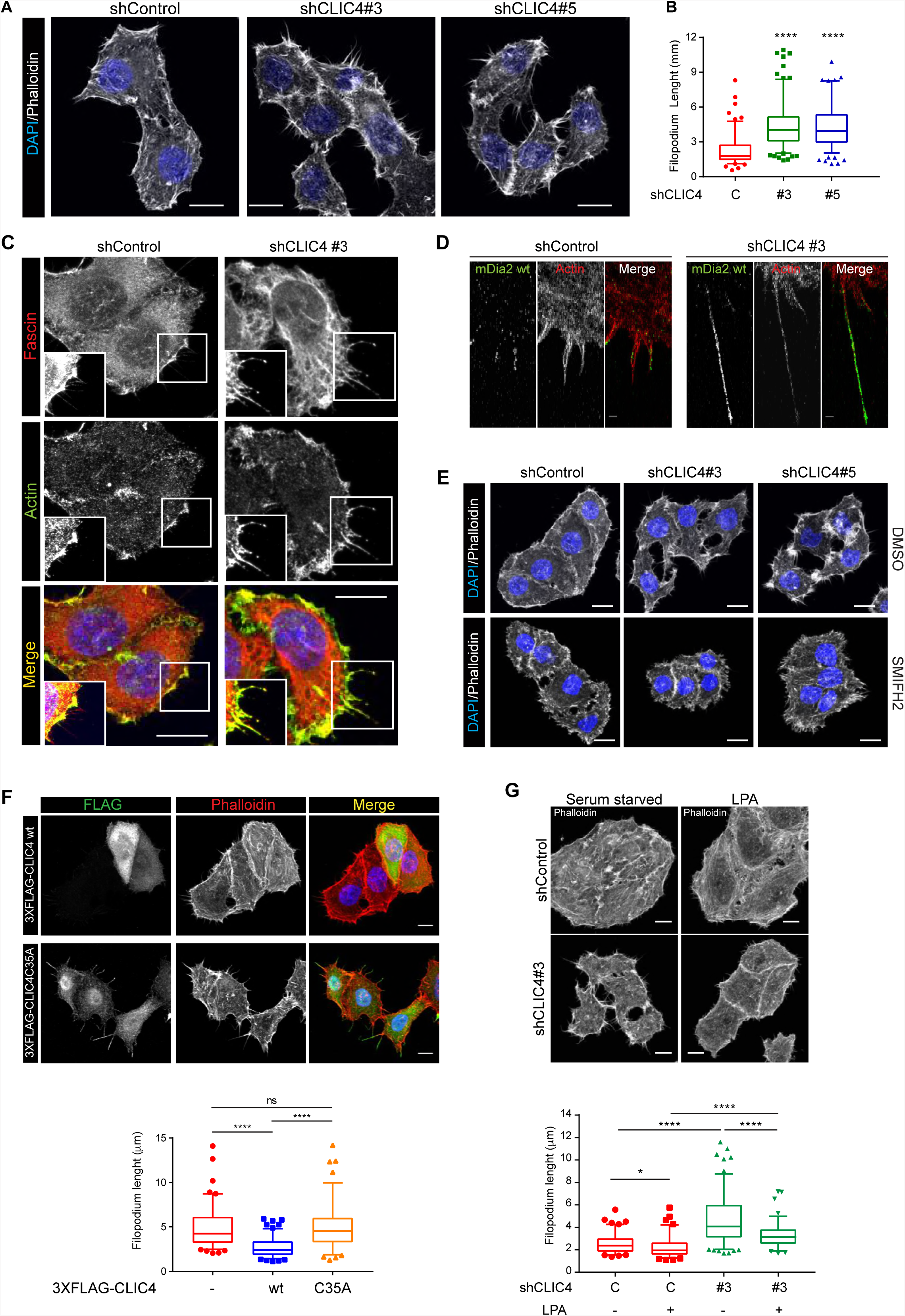
CLIC4 depletion increases filopodium length in HeLa cells. (**A**) Effect of CLIC4 depletion on filopodium lenght. Control and CLIC4 knockdown HeLa cells were seeded on collagen-I-coated glass coverslips and serum starved overnight and fixed. Maximal projections of confocal Z-stacks show actin cytoskeleton and nuclear staining with phalloidin (grey) and DAPI (blue), respectively. Scale bar, 10 µm. (**B**) Quantification of filopodium length. The length of basal filopodia was quantified by using ImageJ software. Data represent filopodium length measured in two independent experiments (n_Control_ = 28, n_shCLIC4#3_ = 34, n_shCLIC4#5_ = 31, n_Rescue_ = 30 cells). (**C**) Fascin staining. Cells were treated as in **A**. Confocal images of cells stained with actin (green), fascin (red) and DAPI (blue) are shown. Scale bar, 10 µm. (**D**) Super Resolution (SR) imaging of mDia2 in filopodia. Cells were seeded on collagen-I-coated glass coverslips, transfected with FLAG-mDia2 wt and serum starved overnight. SR images of mDia2 wt (green) and phalloidin (red). (**E**) SMIFH2 treatment reduces filopodium length. Cells were seeded on collagen-I-coated glass coverslips, serum starved overnight and incubated with SMIFH2 (50 µM, 20 min) or mock-treated (DMSO) before fixation. Maximal projections of confocal Z-stacks show actin cytoskeleton and nuclear staining with phalloidin (grey) and DAPI (blue), respectively. Scale bar, 10 µm. (**F**) Rescue of the CLIC4-deficient phenotype with CLIC4 wt and CLIC4C35A.CLIC4 knockdown HeLa cells were seeded on collagen-I-coated glass coverslips and transfected with a mouse version of CLIC4 wild type (3XFLAG-mCLIC4 wt) and its mutant in Cys35 (3XFLAG-mCLIC4C35A). Cells were serum starved overnight and fixed. Maximal projections of confocal Z-stacks show actin cytoskeleton and nuclear staining with phalloidin (grey) and DAPI (blue), respectively. Scale bar, 10 µm. The length of basal filopodia was quantified by using ImageJ software. Data represent filopodium length measured in two independent experiments (n_non transfectd_=25, n_3XFLAGmCLIC4 wt_= 26, n_3XFLAG-mCLIC4C35A_= 25). (**G**) Effect of LPA stimulation on filopodium length. shControl and shCLIC4 cells were seeded on collagen-I-coated coverslips and serum-starved overnight. Cells were stimulated with LPA (2 µM, 2 min) or left untreated before fixation. Maximal projections of confocal Z-stacks stained with phalloidin (grey) are shown. Scale bar, 10 µm. The length of basal filopodia was quantified as in (**F**). Data represent filopodium length measured in two independent experiments (n_Control starvation_= 25, n_Control LPA_= 24, n_shCLIC4#3 starvation_= 30, n_shCLIC4#3 LPA_= 22 cells). The box represents the 25^th^ to 75^th^ percentiles, and the median is indicated by the line, the whiskers represent the 5^th^and the 95^th^percentiles. ****P<0.001; ns, not significant.

Importantly, this CLIC4 knockdown phenotype could be rescued by reintroducing shRNA-resistant mouse CLIC4 (3XFLAG-mCLIC4) (**Fig. 6F**), whereas the Profilin-1-binding-and trafficking-incompetent CLIC4(C35A) mutant could not. These findings underscore the importance of Profilin-1 binding in dictating the CLIC4-regulated phenotype. Finally, LPA treatment reduced filopodium length in both control and CLIC4-depleted cells (**Fig. 6G**), but filopodia retraction by LPA was less complete upon CLIC4 depletion (**Fig. 6G-H**). Collectively, our findings suggest that recruitment of CLIC4 to the plasma membrane serves to suppress the formation of integrin-regulated filopodia, along with promoting integrin trafficking [10].

### Concluding Remarks

The present study sheds new light on the actin-based functions of CLIC4, in particular its dynamic trafficking to the plasma membrane upon receptor stimulation and the identification of a direct binding partner, namely Profilin-1. Owing to Profilin-1 binding, CLIC4 appears to function in a RhoA-mDia2-regulated actin polymerization network at the cell periphery, where it modulates integrin function, cell adhesion and filopodia (cf. [10]). We have identified distinct residues in CLIC4 that make up a concave surface or elongated “cleft” that binds Profilin-1 and are essential for CLIC4 trafficking upon RhoA activation.

A major challenge for future studies is ti identify Profilin-1 residues that are essential for binding to CLIC4 but do not perturb the interaction with poly-Pro-containing proteins and PIP_2_ [29]. Regardless of the precise CLIC4-Profilin-1 binding mode, our studies support a model in which RhoA-mDia2-driven actin polymerization triggers trafficking of the CLIC4-PFN1 complex to the plasma membrane to integrate actin dynamics, integrin trafficking and filopodia protrusion. CLIC4 has.also been implicated in inhibiting the formation of branched actin on early endosomes through an unknown mechanism [13]. Binding of CLIC4 to Profilin-1 and the ability of Profilin-1 to attenuate branched actin formation [36] may thus explain this puzzling observation. We therefore propose that Profilin-1 could be a general modulator of CLIC4 actin-dependent processes. Profilin-1 is known to interact with multiple ligands and therefore is implicated in signaling processes beyond its actin-binding properties; furthermore, several studies have recently emerged that link mutations in *PFN1* to various human diseases [37,38]. Since some of these *PFN1* mutations are located in the poly-Pro binding side, it will be interesting to examine their potential effect on CLIC4 function. Finally, future studies should assess whether the functions of other CLIC family members similarly rely on Profilin-1 binding.

## Materials and Methods

### Reagents and antibodies

1-oleoyl-LPA, Y-27632 and SMIFH2 were from Sigma. EGF was from PeproTech. Type-I collagen was from Inamed BioMaterials. EDTA-free protease inhibitor cocktail tablets were from Roche. Phalloidin-red (acti-stain™555 Phalloidin) was from Cytoskeleton, Inc.. SMIFH2 was used as previously described [19] Antibodies used were: monoclonal anti-β-actin (AC-15) and anti-FLAG M2, Sigma; polyclonal anti-CLIC4, anti-mDia2 and anti-GFP, generated in-house [10,19]; monoclonal anti-mDia1 (D3) and polyclonal anti-Profilin1, Santa Cruz; Alexa-Fluor-conjugated secondary antibodies (Invitrogen).

### Vectors

pGEX-6P1-mDia2 (FH1-FH2), pCDNA3 Flag-mDia2 wt, pGEX-6P1-hPFN1 and pEGFP-C1-hPFN1 were previously described [18,33,39]. pGEX-6P1-CLIC4 wt was previously described [10]. pGEX-6P1-CLIC4(C35A) was generated from pGEX-6P1-CLIC4 wt using Phusion Site-Directed Mutagenesis Kit (Thermo Fisher Scientific). For rescue experiments, 3XFLAG-tagged mouse CLIC4 wt (mCLIC4) and CLIC4(C35A) were cloned in LZRS-Blast vector. LZRS-IRES-GFP-RhoA(N19) and LZRS-IRES-GFP-RhoA(V14) were a gift from Dr. Jacques Neefjes.

### Cell Culture, infections and transfections

HeLa cells were grown in Dulbecco’s modified Eagle’s medium (DMEM, Invitrogen) supplemented with 10% fetal bovine serum (FBS) under 5% CO_2_ at 37 °C. Stable HeLa CLIC4 and mDia1 knockdown cells were previously described and characterized [10,18] [21]. For mDia2 and Profilin1 knockdown studies, four distinct shRNAs in the lentiviral vector pLKO.1 were employed (TRC human shRNA library; Sigma TRCN0000150903 (shmDia2#817) and TRCN0000150850 (shmDia2#820) [18], TRCN0000311689 (shProfilin1#1), TRCN0000294209 (shProfilin1#3)). pLKO.1 empty vector was used as control. Lentiviral production and stable cell infection were performed as previously described [10]. Plasmid transfections for imaging studies were performed with X-tremeGene 9 (Roche) reagent according to manufacturer’s instructions.

### Live cell imaging, immunofluorescence and image analysis

For live cell imaging experiments, cells were seeded on 24 mm glass coverslips and transiently transfected with YFP-CLIC4 wild type or C35A mutant. Cells were serum starved overnight with DMEM supplemented with 0.1% FBS (starvation medium) and imaged in DMEM-F12 without Phenol Red (Invitrogen) under 5% CO_2_ at 37 °C on a Leica TCS-SP5 confocal microscope (63X objective). We used minimal laser intensity to avoid photobleaching and phototoxicity. Series of confocal images were taken at 5 seconds. LPA was added 2 minutes after starting imaging. The translocation of YFP-CLIC4 was quantified using ImageJ software. Confocal imaging of fixed cells was performed as previously described [10]. Filopodium length at the basal membrane of phalloidin-stained cells was measures manually using ImageJ software. Protrusions shorter than 2 µm were not considered for quantification.

### Super-resolution imaging

For super-resolution microscopy using the ground-state depletion imaging method [40], cells were cultured on 24 mm, no. 1.5 coverslips and transiently transfected with FLAG-mDia2. After 24 h, cells were serum starved overnight, washed briefly with PBS, fixed with 4% PFA for 10 min at room temperature and permeabilized with 0.1% Triton X-100. Samples were extensively washed with PBS and blocked with 5% BSA for 30 min at room temperature. Coverslips were incubated for 1 h with anti-FLAG M2 (Sigma) primary antibody at room temperature, washed and incubated with Alexa-Fluor-conjugated secondary antibody (Alexa Fluor 647 and 488) and Phalloidin (Alexa Fluor 488 and 647). Super-resolution microscopy was performed with a Leica SR GSD microscope (Leica Microsystems, Wetzlar, Germany) mounted on a Sumo Stage (#11888963) for drift free imaging. Collection of images was done with an EMCCD Andor iXon camera (Andor Technology, Belfast, UK) and an oil immersion objective (HCX PL Apo 100X, NA 1.47). Laser characteristics were 405nm/30mW, 488 nm/300mW and 647nm/500mW, with the 405nm laser used for back pumping and the others for wide field/TIRF imaging. Ultra clean coverslips (cleaned and washed with base and acid overnight) were used for imaging. The number of recorded frames was variable between 10,000 to 50,000, with a frame rate of 100Hz. The data sets were analyzed with the Thunder Storm analysis module [41] and images were reconstructed with a detection threshold of 70 photons, sub pixel localization of molecules and uncertainty correction, with a pixel size of 10 nm.

### Western Blotting

Whole-cell lysates were prepared by lysing cells in JS lysis buffer (50 mM HEPES, pH 7.5, 150 mM NaCl, 1.5 mM MgCl_2_, 5 mM EGTA, 1% glycerol, 0.5% Triton X-100) supplemented with NaO_3_V_4_ (5 µM), NaF (1 µM), and protease inhibitor cocktail (Roche). Membranes were blocked in non-fat dry milk and incubated with primary antibodies according to the manufacturers’ instructions, followed by HRP-conjugated secondary antibodies (Dako Inc.,1:10,000).

### Recombinant proteins

GST-tagged CLIC4 (wt and C35A) was expressed in BL21 E. coli and CLIC4 proteins were purified using Glutathione Sepharose^®^ 4B beads (GE Healthcare) and gel filtration. GST-tag was removed with PreScission Protease (GE Healthcare). Expression, purification and cleavage of human Profilin-1 were performed as previously described [39]. Expression and purification of mDia2 (FH1-FH2 domain) were previously described [33], whereas cleavage done as described for Profilin-1. Expression and purification of Rho-GTPase binding domains was previously described [42].

### FRET-based RhoA Biosensor

The design and use of the FRET-based RhoA biosensor was described previously [17]. Briefly, the complete amino acid sequence of a RhoA was positioned at the C terminus of a single polypeptide chain to preserve its interaction with GDI and other regulatory proteins. A FRET pair consisting of Cerulean3 and circularly permutated Venus was used. The HR1 region of PKN was used as the effector domain for activated RhoA. In a control biosensor, point mutation L59Q in PKN was introduced to generate a binding-deficient effector domain, so that FRET ratios remained at the basal level regardless of the activation state of RhoA.

### RhoA and Cdc42 pull-down assays

Rho-GTPase pull-down assays were performed as previously described [33,39].

### Actin polymerization assays

Actin purification, bulk actin polymerization assays and F-actin co-sedimentation assays were performed as previously described [43,44].

### Molecular modeling

We used the CLIC4 structure from PDB 2AHE and the Profilin-1 structure from PDB 2PAV. For the HADDOCK modeling, we used the web interface in the expert mode. For CLIC4 we defined F37, P76, D97 and Y244A as “active” residues involved in binding. For Profilin-1 we extracted all the residues that are in the surface but do not interact with actin in the 2PAV structure using AREIMOL from the CCP4 suite [45]. 10,000 models were generated in HADDOCK, of which 500 were refined and the best 200 were refined with waters and clustered. 196 out of the final water-refined 200 models clustered in four clusters with 132, 45, 10, and 9 models which respectively had HADDOCK scores of −122, −89, −90, and −87 and Z-scores of −1.7, 0.6, 0.5 and 0.7 respectively. The top cluster also showed a low internal RMSD (0.7 +/- 0.5) and was clearly representing the most likely model for the interaction. Details of the HADDOCK scoring are shown in **Suppl. Table 1**. Molecular structures in Fig. 5 were prepared by CCP4MG [46].

### Statistical analysis

For determination of statistical significance, unpaired Student’s t-tests were performed using GraphPad Prism software. The indicated significance values are compared to control conditions. P-values are indicated as follows: *P<0.05, **P<0.01, ***P<0.001, ****P<0.0001.

## Acknowledgements

We thank Tatjana Heidebrecht for providing recombinant proteins for biophysical experiments performed by Alexander Fish at the NKI Protein Facility, and Alexandre Bonvin for advice using HADDOCK. The FP7 WeNMR (project# 261572) and H2020 West-Life (project# 675858) European e-Infrastructure projects are acknowledged for the use of their web portals, which make use of the EGI infrastructure and the DIRAC4EGI service with the dedicated support of CESNET-MetaCloud, INFN-PADOVA, NCG-INGRID-PT, RAL-LCG2, TW-NCHC, IFCA-LCG2, SURFsara and NIKHEF, and the additional support of the national GRID Initiatives of Belgium, France, Italy, Germany, the Netherlands, Poland, Portugal, Spain, UK, South Africa, Malaysia, Taiwan and the US Open Science Grid.

## Author contributions

EA, WHM and MI initiated the research; EA, KK, LN, TI, KJ, AP and MI developed methodology; EA, KK, LN, TI and MI performed experiments and collected data; AP, performed molecular modelling; LN, performed super-resolution microscopy; KJ, supervised microscopy and live-imaging studies; EA, KK, LN, KJ, AP, WHM and MI analysed and interpreted data; EA, WHM and MI wrote the manuscript with input from all authors.

## SUPPLEMENTAL FIGURES

**Suppl. Figure 1.**
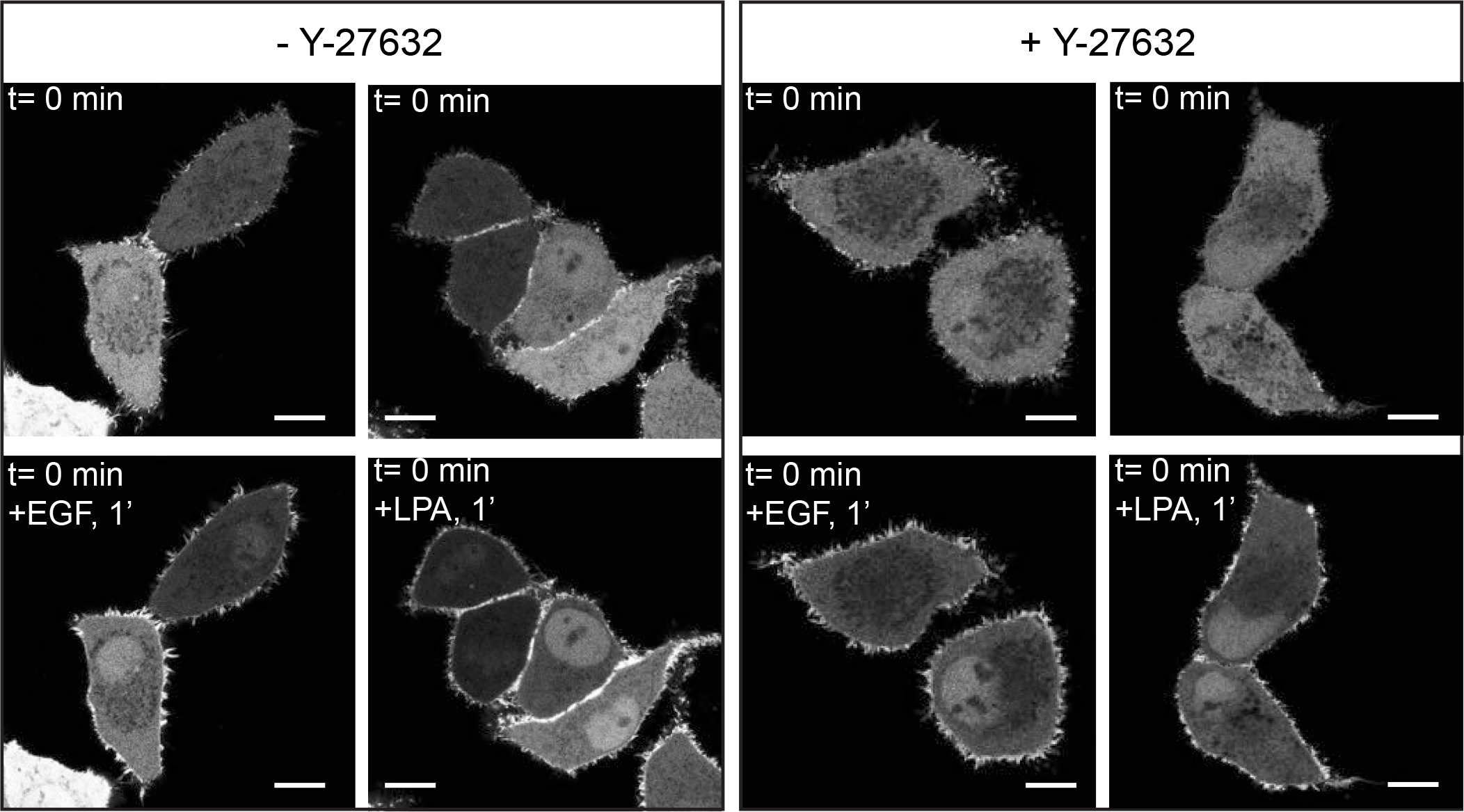
LPA- and EGF-induced translocation of CLIC4 does not depend on ROCK activation. CLIC4 translocation does not depend on ROCK. Cells were transfected with YFP-CLIC4 and pre-incubated with Y-27632 (10 µM, 30 minutes) before LPA (2 µM) or EGF (100 ng/ml) stimulation. DMSO-treated cells were used as control. Frames from time-lapse movies at the indicated time points are shown. Scale bar: 10 µm.

**Suppl. Figure 2.**
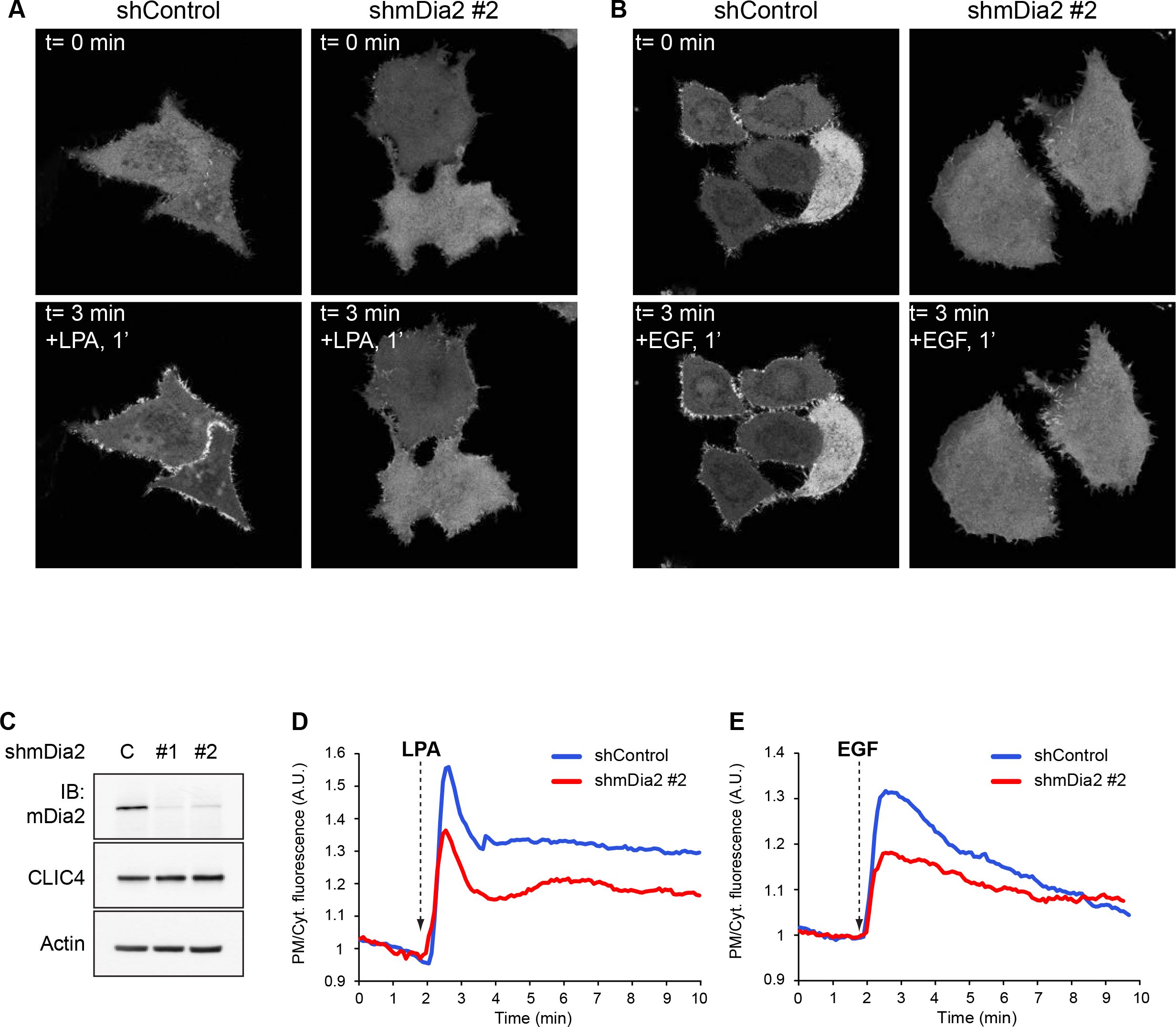
LPA-induced translocation of CLIC4 depends on mDia2. (**A**) Live-cell imaging of CLIC4 translocation in mDia2 knockdown HeLa cells. A stable population of mDia2 knockdown cells using an independent hairpin (shmDia2#820) was obtained as described in the Materials and Methods. shControl, shmDia2 cells were seeded on glass coverslips and transfected with YFP-CLIC4. LPA (2 µM) was added 2 minutes after starting imaging. SMIFH2 was maintained during LPA stimulation Frames from time-lapse movies at the indicated time points are shown. Scale bar: 10 µm. (**B**) mDia2 knockdown efficiency was determined by immunoblotting (IB) using anti-mDia2 antibody β-actin was used as loading control. (C) Quantification of CLIC4 translocation. Net translocation is expressed as PM/Cyt fluorescence ratio (n_shcontrol_= 11, n_shmDia2#820_= 14, from two independent experiments).

**Suppl. Figure 3.**
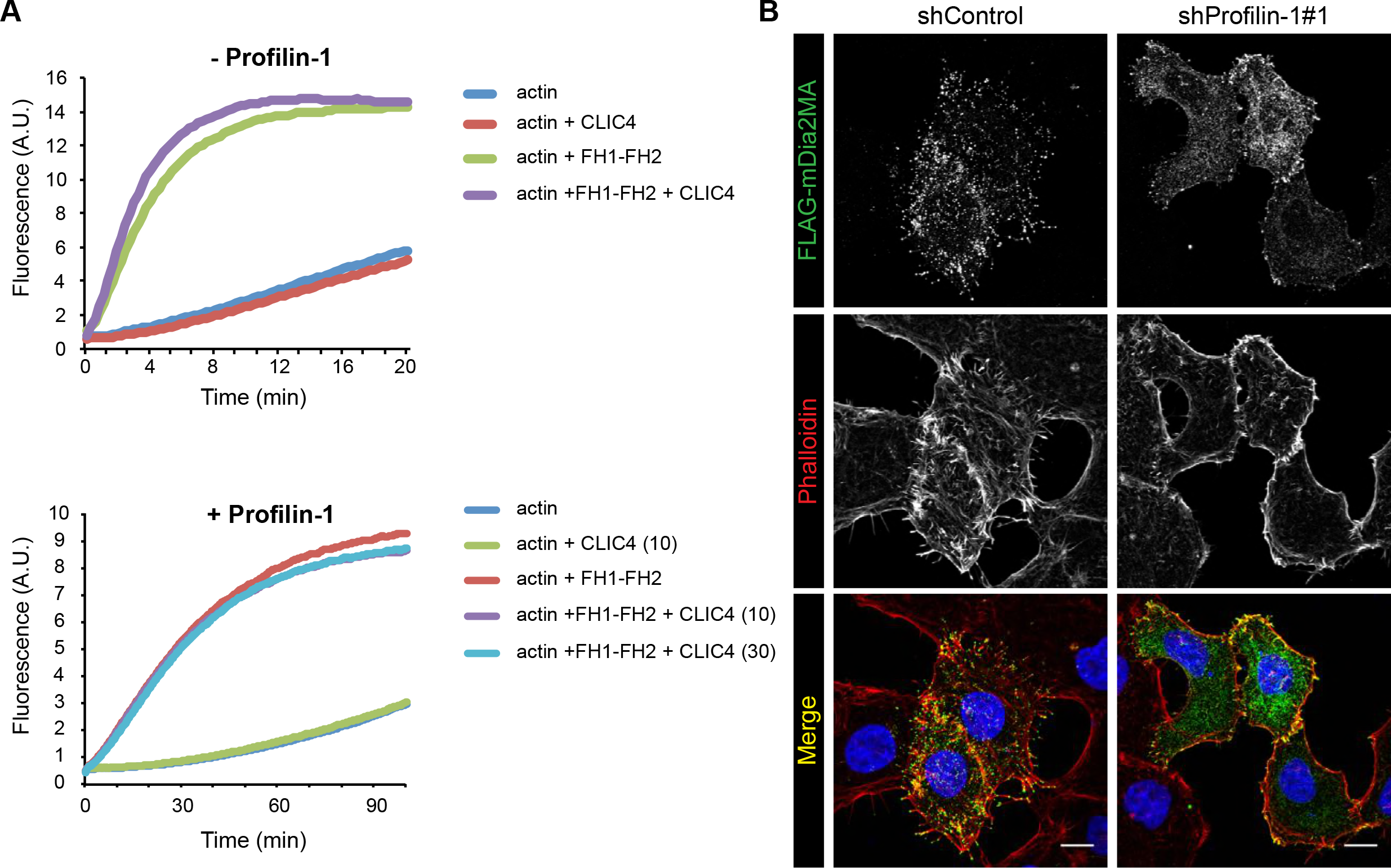
Actin polymerization *in vitro* and characterization of Profilin-1-depleted cells. (**A**) Actin polymerization assay as described in Material and Methods using purified recombinant CLIC4 (10 µM and 30 µM), actin (1-2 µM), mDia2-FH1-FH2 (0.1 µM) without (top) or with (bottom) Profilin-1 (5 µM). (**B**) Profilin-1 knockdown cells do not form filopodia. Control and CLIC4 knockdown HeLa cells were seeded on collagen-I-coated glass coverslips and transfected with constitutively active mDia2 (FLAG-mDia2MA). Cells were serum starved overnight and fixed. Maximal projections of confocal Z-stacks show phalloidin (red), mDia2MA (green) and DAPI (blue). Scale bar, 10 µm.

## Supplemental Movies

**Movie 1. CLIC4 translocation induced by LPA in HeLa cells.** Note the transient character of CLIC4 accumulation at the plasma membrane, concomitant with homogenous depletion of cytosolic GFP-CLIC4 upon addition of LPA (1 µM). Images were collected every 10 seconds and shown at 5 frames per second.

**Movie 2. CLIC4 translocation induced by EGF in HeLa cells.** Experimental conditions as in Movie 1. EGF was added at 100 nM.

**SUPPLEMENTAL TABLE 1.** HADDOCK modeling parameters.

|  | **Cluster 1** | **Cluster 2** | **Cluster 3** | **Cluster 4** |
| --- | --- | --- | --- | --- |
| **HADDOCK score** | -121.8 ± 2.8 | -89. ±2.0 | -90.8 ±6.7 | -87.0 ± 4.5 |
| **Cluster size** | 132 | 45 | 10 | 9 |
| **RMSD** | 0.7 ± 0.5 | 13.0 ± 0.0 | 9.9 ± 0.2 | 9.2 ±0.6 |
| **Van der Waals energy** | -58.9 ± 5.2 | -49.3 ± 8.3 | -52.4 ± 6.1 | -52.0 +/- 5.2 |
| **Electrostatic energy** | -291.5 ±54.1 | -228.3 ± 51.2 | -226.6 ± 24.8 | -177.9 ±<br>33.8 |
| **Desolvation energy** | -4.7 ± 6.2 | 5.5 ± 9.9 | 6.8 ± 12.1 | 0.4 +/- 6.5 |
| **Restraints violation e.** | 1.2 ± 0.78 | 2.7 ± 3.12 | 1.3 ± 0.79 | 0.8 +/- 0.40 |
| **Buried Surface Area** | 1979.0 ± 65.8 | 1702.0 ± 105.3 | 1613.1 ± 82.5 | 1656.7 ±<br>58.3 |
| **Z-Score** | -1.7 | 0.6 | 0.5 | 1656.7 ±<br>58.3 |

